# Stimulus prediction in the hippocampus

**DOI:** 10.1101/246843

**Authors:** Peter Kok, Nicholas B. Turk-Browne

**Affiliations:** Princeton Neuroscience Institute and Department of Psychology, Princeton University, Princeton, NJ 08544; Department of Psychology, Yale University, 2 Hillhouse Avenue, New Haven, CT 06520

**Keywords:** Expectation, perceptual inference, predictive coding, shape perception, hippocampal subfields

## Abstract

Perception can be cast as a process of inference, in which bottom-up signals are combined with top-down predictions in sensory systems. However, the source of these top-down predictions, especially when complex and multisensory, remains largely unknown. We hypothesised that the hippocampus — which rapidly learns arbitrary associations and has bidirectional connections with sensory systems — may be involved. We exposed humans to auditory cues predicting visual shapes, while measuring high-resolution fMRI signals in visual cortex and the hippocampus. Using multivariate reconstruction methods, we discovered a dissociation between these regions: representations in visual cortex were dominated by whichever shape was presented, whereas representations in the hippocampus (CA3 and subiculum, but not CA1) reflected only which shape was predicted by the cue. The strength of hippocampal predictions correlated across participants with the amount of expectation-related facilitation in visual cortex. These findings are consistent with the possibility that the hippocampus supplies predictions to sensory systems.

## Introduction

Neural activity in sensory cortex can be strongly modulated by prior expectations (Summerfield et al., 2008; Den Ouden et al., 2009; Alink et al., 2010; Meyer and Olson, 2011; Todorovic et al., 2011; Wacongne et al., 2011; Kok et al., 2012, 2013; Bell et al., 2016; Kaposvari et al., 2016). However, the source of such top-down predictions remains unknown. Models of predictive coding emphasize the role of downstream areas within local brain circuits (Rao and Ballard, 1999; Spratling, 2010). Although this could account for basic, highly ingrained predictions, such as surround suppression or filling-in of contours (Lee and Nguyen, 2001; Spratling, 2010; Kok and De Lange, 2014), it is unclear how this mechanism applies to more complex, learned predictions. Consider cross-modal predictions, for example, such as when an auditory stimulus (e.g., a bell or bark) leads to an expectation of the visual appearance of the corresponding object (e.g., a bicycle or dog). Such associations, especially when learned recently, cannot readily be encoded within sensory systems, as visual cortex does not have direct access to the features of auditory stimuli nor is it able to rapidly bind these features. Such predictions may instead depend on a higher-order brain region that can learn multisensory associations in the world, retrieve them based on partial information (e.g., a sound), and reinstate missing information (e.g., the associated visual object) in relevant sensory cortex.

Based on these desiderata, we hypothesised that the hippocampus plays a role in such predictions. First, the hippocampus is known to be involved in learning associations between arbitrary stimuli (Cohen and Eichenbaum, 1993; Davachi, 2006; Turk-Browne et al., 2009; Henke, 2010; Hsieh et al., 2014; Garvert et al., 2017), particularly when these stimuli are discontiguous in time or space (Wallenstein et al., 1998; Staresina and Davachi, 2009). In fact, learning of such relationships is strongly impaired when the hippocampus is damaged (Sutherland et al., 1989; Chun and Phelps, 1999; Hannula et al., 2006; Konkel et al., 2008; Schapiro et al., 2014). Second, the hippocampus has bidirectional connections with sensory cortices of all modalities (Lavenex and Amaral, 2000; Eichenbaum et al., 2007; Henke, 2010), and has for this reason even been considered the top of sensory hierarchies (Felleman and Van Essen, 1991). Third, one of the main computational functions of the hippocampus is to retrieve associated items from memory based on partial information, a process known as pattern completion (Treves and Rolls, 1994; McClelland et al., 1995; Henke, 2010). This function has been mostly considered in the context of recall from episodic memory, but is also ideally suited for retrieving predictions based on contextual cues (McClelland et al., 1995; Eichenbaum and Fortin, 2009; Schapiro et al., 2012; Davachi and DuBrow, 2015; Hindy et al., 2016). Pattern completion is thought to be subserved by the CA3 subfield of the hippocampus, because of its strong recurrent, autoassociative connections (Treves and Rolls, 1994; Henke, 2010; Schapiro et al., 2017), from whence the retrieved pattern is sent to CA1, where it may be compared to actual sensory inputs (Lisman and Grace, 2005; Chen et al., 2011; Duncan et al., 2012). Additionally, this information can be fed back to sensory cortex, including via the subiculum, with retrieved memories reinstated in the cortical areas that initially encoded them during perception (McClelland et al., 1995; Bosch et al., 2014; Gordon et al., 2014; Rothschild et al., 2017). Again, this cortical reinstatement has been mainly considered in the context of episodic memory retrieval (Davachi and Danker, 2013), but may also subserve prediction, especially when a contextual cue precedes the associated item (McClelland et al., 1995; Eichenbaum and Fortin, 2009; Kok et al., 2014; Reddy et al., 2015; Hindy et al., 2016; Kok et al., 2017).

To investigate the involvement of the hippocampus in cross-modal predictions, we exposed human participants to auditory tones preceding the appearance of particular visual shapes (Figure 1), while measuring signals in both visual cortex and the hippocampus with high-resolution functional magnetic resonance imaging (fMRI). Using multivariate pattern analysis to reconstruct shapes from BOLD activity (Figure 2), we explored what visual information was represented in these brain systems on trials in which the tones validly vs. invalidly predicted which shape would appear. We hypothesised that the hippocampus would represent the expected shape regardless of what appeared. In contrast, visual cortex may be modulated by expectation, but its representation should be dominated by the shape that was presented on screen. Finally, we hypothesised that there would be a relationship between hippocampal predictions and effects of prediction on visual cortex, which would be consistent with a role for the hippocampus in supplying sensory expectations.

**Figure 1.**
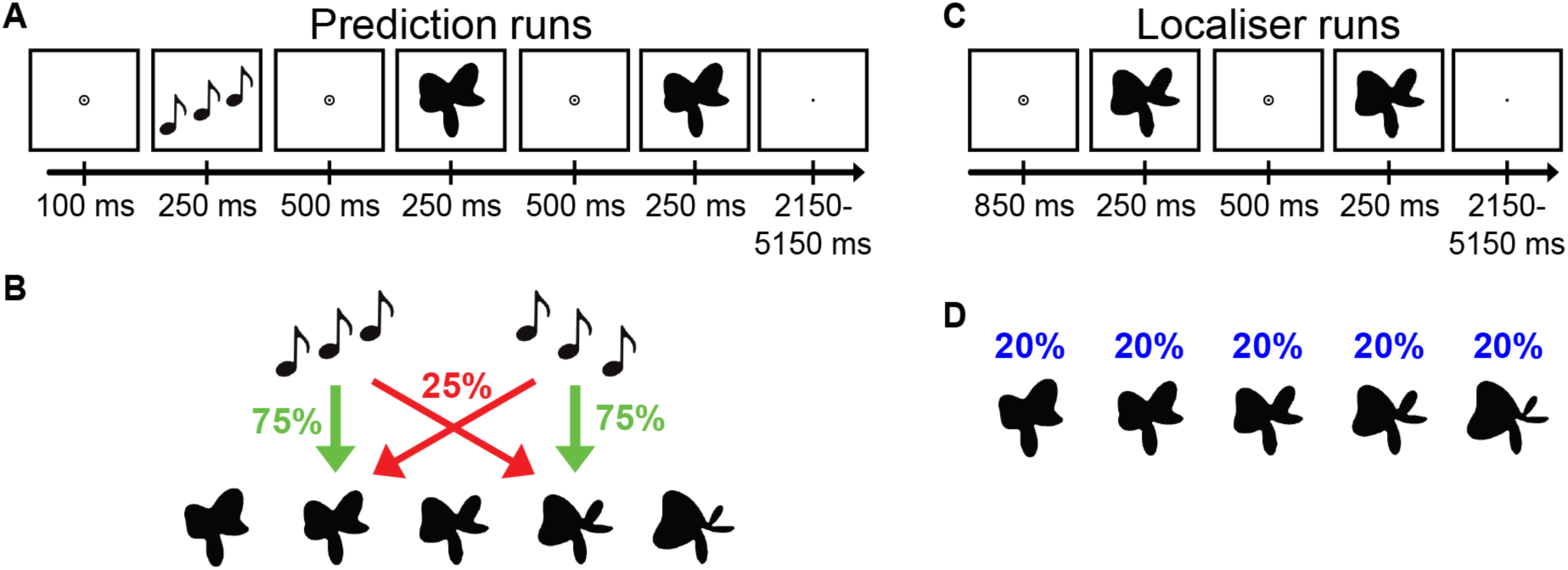
Experimental paradigm. (**A**) During prediction runs, an auditory cue preceded the presentation of two consecutive shape stimuli. On each trial, the second shape was either identical to the first or slightly warped with respect to the first along an orthogonal dimension, and participants’ task was to report whether the two shapes were the same or different. (**B**) The auditory cue (ascending vs. descending tones) predicted whether the first shape on that trial would be shape 2 or shape 4 (out of five shapes). The cue was valid on 75% of trials, while in the other 25% of (invalid) trials the unpredicted shape was presented. (**C**) During localiser runs, no auditory cues were presented. As in the prediction runs, two shapes were presented on each trial, and participants’ task was to report same or different. (**D**) All five shapes appeared with equal (20%) likelihood on trials of the localiser runs.

**Figure 2.**
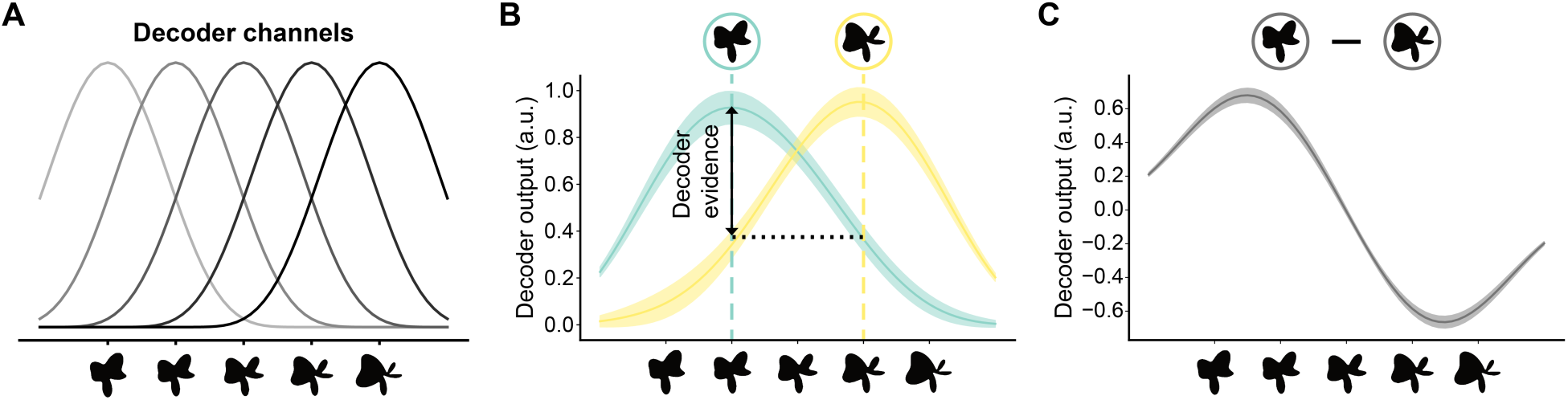
Illustration of the decoding method. (**A**) We used a forward modelling approach to reconstruct shapes from the pattern of BOLD activity. Shape selectivity was characterised by five hypothetical channels, each with an idealised shape tuning curve. BOLD patterns obtained from the localiser runs were used to estimate the weights on the five hypothetical channels separately for each voxel, using linear regression. (**B**) Using these weights, the second stage of the analysis reconstructed the channel outputs associated with the pattern of activity across voxels evoked by the prediction runs (only shapes 2 and 4 were used in these runs). Channel outputs were converted to a weighted average of the five basis functions, resulting in neural shape tuning curves. Decoding performance was quantified by subtracting the amplitude of the shape tuning curve at the presented shape (e.g., shape 2) from the amplitude at the non-presented shape (e.g., shape 4). (**C**) Finally, we collapsed across the presented shapes by subtracting the shape tuning curve for shape 4 from that for shape 2, thereby removing any non-shape-specific BOLD signals. Shaded regions in **B** and **C** indicate SEM.

## Results

Participants were exposed to auditory tones that validly or invalidly predicted the upcoming shape stimulus (Figure 1A-B). This first shape was followed by a second shape that was either identical to the first, or slightly warped. Participants performed a shape discrimination task, reporting whether the two shapes on a given trial were the same or different.

### Behaviour

Participants were able to discriminate small differences in the complex shapes, during the localiser runs (36.9% ± 2.3% modulation of the 3.18 Hz radial frequency component, mean ± SEM) and during the prediction runs (valid trials: 31.6% ± 2.5%; invalid trials: 33.2% ± 2.9% modulation). The discrimination thresholds for valid and invalid trials were not reliably different (*t*_23_ = 1.00, *p* = 0.32). This is not surprising, as the discrimination task was independent of the prediction manipulation: the auditory cue provided no information about which choice was correct and the shape manipulation on different trials was orthogonal to the feature dimensions defining the shape space. Accuracy and reaction times also did not differ significantly between valid (accuracy: 70.6% ± 1.2%; RT: 575 ms ± 16 ms) and invalid trials (accuracy: 68.8% ± 1.5%; RT: 573 ms ± 18 ms; both *p*s > 0.20), which was expected because these conditions were staircased separately to the same performance level.

### Shape reconstruction

The decoder successfully reconstructed the presented shapes from the pattern of activity in visual cortex (V1: *t*_23_ = 14.72, *p* = 3.4 × 10^−13^; V2: *t*_23_ = 14.23, *p* = 6.8 × 10^−13^; LO: *t*_23_ = 7.04, *p* = 3.5 × 10^−7^), with a modest but significant modulation by the predictive cues in V1 (*t*_23_ = 2.58, *p* = 0.017; Figure 3A) but not in V2 (*t*_23_ = 1.42, *p* = 0.17) or LO (*t*_23_ = 0.17, *p* = 0.87). In other words, shape representations in visual cortex were dominated by what was presented to the eyes.

The results were strikingly different in the hippocampus. Here, the pattern of activity contained a representation of the *predicted* shape (*t*_23_ = 2.86, *p* = 0.0089), while the presented shape was not significantly represented (*t*_23_ = 0.54, *p* = 0.59). That is, shape representations in the hippocampus were fully determined by the auditory cue and the expectation it established (Figure 3B).

**Figure 3.**
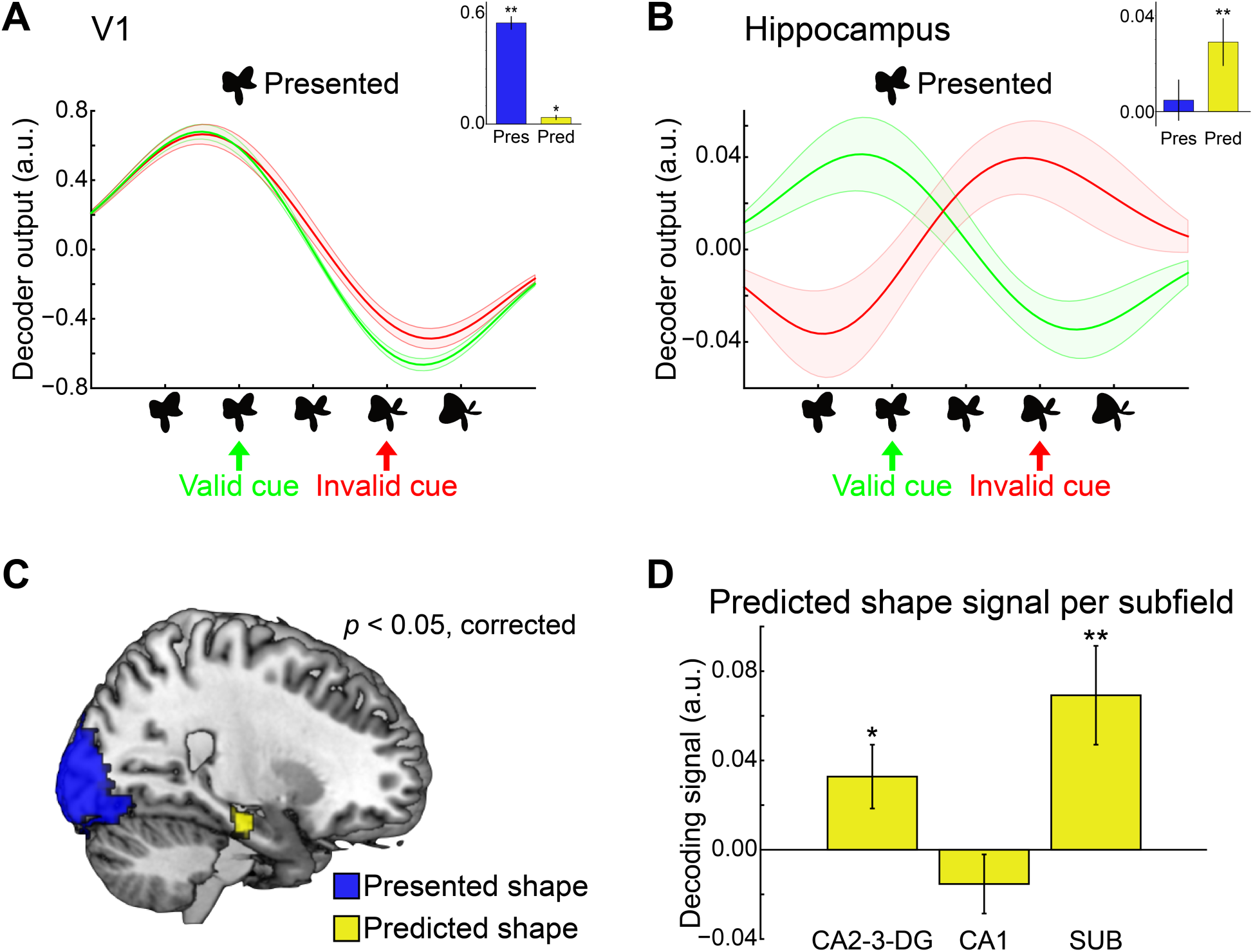
Shape representations in visual cortex and hippocampus. (**A**) Shape reconstructions from patterns of activity in V1, separately for validly (green) and invalidly (red) predicted shapes. Representations in visual cortex (V1, V2, LO; V1 plotted as representative region) were dominated by the presented shape, with modest modulation by the predictive cues in V1. Inset depicts quantified evidence for presented (blue) and predicted (yellow) shapes. (**B**) Shape reconstructions in the hippocampus were fully determined by the cued (predicted) shape, rather than the presented shape. Inset depicts quantified evidence for presented (blue) and predicted (yellow) shapes. (**C**) A searchlight analysis revealed evidence for the presented shape in the occipital lobe and for the predicted (but not presented) shape in the hippocampus. See Table 1 for full results. (**D**) Decoding of the predicted shapes across hippocampal subfields. \**p* < 0.05, \*\**p* < 0.01. Shaded regions and error bars indicate SEM.

This dissociation was confirmed by a searchlight analysis, which revealed significant evidence for the presented shape in the occipital lobe and for the predicted (but not presented) shape in the hippocampus (Figure 3C). This analysis also revealed evidence for the predicted shape in more anterior occipital cortex and a few smaller clusters elsewhere (Table 1).

**Table 1.** Searchlight results

| Anatomical region | Hemisphere | Cluster size | Peak $p$ | Coordinates (x y z) | | |
| --- | --- | --- | --- | --- | --- | --- |
| <u>Presented shape decoding</u> |  |  |  |  |  |  |
| Posterior occipital cortex | Bilateral | 8964 | < 0.001 | -22 | -88 | -22 |
| <u>Predicted shape decoding</u> |  |  |  |  |  |  |
| Calcarine sulcus | Right | 180 | 0.016 | 14 | -70 | 20 |
| Hippocampus | Right | 63 | 0.026 | 24 | -18 | 16 |
| Middle cingulate | Right | 27 | 0.028 | 2 | 8 | 40 |
| Caudate | Left | 7 | 0.028 | -16 | 4 | 8 |
| Cerebellum | Left | 5 | 0.044 | -26 | -62 | 22 |
| All $p$-values are corrected for multiple comparisons. Coordinates reflect local maxima of significant clusters in Montreal Neurological Institute (MNI) space. | | | | | | |

Note that the hippocampal cluster in the searchlight analysis was in the right hemisphere only, whereas we collapsed over hemisphere in the ROI analysis. This apparent laterality may be an artefact of statistical thresholding or may indicate a genuine hemispheric difference. We did not have hypotheses about left vs. right or anterior vs. posterior hippocampus, but investigated these divisions *post hoc* because of the searchlight results by subdividing the hippocampal ROI. There were no reliable differences in evidence for the predicted shape in left vs. right (*p* = 0.72) or anterior vs. posterior (*p* = 0.93) hippocampus. In fact, decoding of the predicted shape was significant within each of these four subdivisions of the hippocampus individually (all *p*s < 0.05).

To investigate the circuitry underlying these predictions further, we applied an automated anatomical segmentation method to distinguish the subfields of the hippocampus. This analysis revealed that hippocampal subfields encoded the predicted shapes to different extents (*F*_2,46_ = 6.45, *p* = 0.0034; Figure 3D). The CA3 subfield is thought to be most strongly involved in pattern completion — i.e., retrieving previously encoded memories from partial cues — whereas CA1 compares such retrieved memories to incoming sensory input supplied by entorhinal cortex (EC). Accordingly, we hypothesised that CA3 would have a purer representation of the predicted shape than CA1, since the latter would also be affected by the presented shape. In line with this hypothesis, predicted shapes could be reconstructed in CA3 (combined with CA2 and dentate gyrus; *t*_23_ = 2.04, *p* = 0.053), and better than in CA1 (*t*_23_ = -1.08, *p* = 0.29; difference between ROIs: *t*_23_ = 2.31, *p* = 0.031). Surprisingly, predicted shapes were also strongly represented in the subiculum (*t*_23_ = 2.97, *p* = 0.0069), which could perhaps be related to its known role in relaying hippocampal signals back to sensory cortex.

We interpret the presence of shape expectations in the hippocampus as reflecting relational memory (Cohen and Eichenbaum, 1993): item memories of the tones and shapes are bound together in a temporal relation during the practice phase and then further during valid trials; when a particular tone cue is encountered, its item memory retrieves this relation and reactivates the item memory for the associated shape. This framework suggests that the success of our decoder depends on the extent to which it has learned about the item memories for different shapes. We examined this hypothesis by breaking down our training examples based on familiarity with the shapes, separating the two localiser runs rather than collapsing, as was done in all of the analyses above. Specifically, we anticipated that training the decoder on the second localiser run at the end of the session (run 6), after participants had the opportunity to repeatedly encode the shapes, would be more effective than training on the first localiser run at the beginning of the session (run 1), when the shapes were more novel. Indeed, we found a significantly stronger representation of the predicted shape when the reconstruction model was trained on the last vs. first localiser run in CA2-CA3-DG (*t*_23_ = 3.09, *p* = 0.0052), but not in CA1 (*t*_23_ = −0.23, *p* = 0.82) or the subiculum (*t*_23_ = 0.89, *p* = 0.38). In visual cortex, on the other hand, training on the last vs. first localiser run did not affect the representation of the predicted shape (V1: *t*_23_ = −0.88, *p* = 0.39; V2: *t*_23_ = 0.17, *p* = 0.87; LO: *t*_23_ = 0.004, *p* = 0.997).

### Visual facilitation

As reported above, shape representations in visual cortex were dominated by the shapes presented to the eyes. However, the temporal evolution of these representations was strongly affected by the auditory prediction cues. We characterised the timecourses of both the mean BOLD response and the shape decoding signal by fitting a canonical (double-gamma) hemodynamic response function (HRF) and its temporal derivative. The parameter estimate of the canonical HRF indicates the peak amplitude of the signal, whereas the temporal derivative parameter estimate reflects the latency of the signal (Friston et al., 1998; Henson et al., 2002).

This approach revealed that there was a modest but highly reliable difference in the latency of the BOLD response evoked by validly and invalidly predicted shapes, as measured by the temporal derivative (V1: *t*_23_ = 6.33, *p* = 1.9 × 10^−6^; V2: *t*_23_ = 7.31, *p* = 1.9 × 10^−7^; LO: *t*_23_ = 7.48, *p* = 1.32 × 10^−7^; Figure 4A). In other words, the BOLD response in visual cortex was significantly delayed by invalid auditory cues. Note that this was not caused by the stimuli *per se*, as the tone-shape mappings were arbitrary and reversed halfway through the study. There was no significant difference in the amplitude of the BOLD response between conditions, as measured by the canonical HRF (V1: *t*_23_ = 0.84, *p* = 0.41; V2: *t*_23_ = 1.64, p = 0.12; LO: *t*_23_ = 1.96, *p* = 0.063).

**Figure 4.**
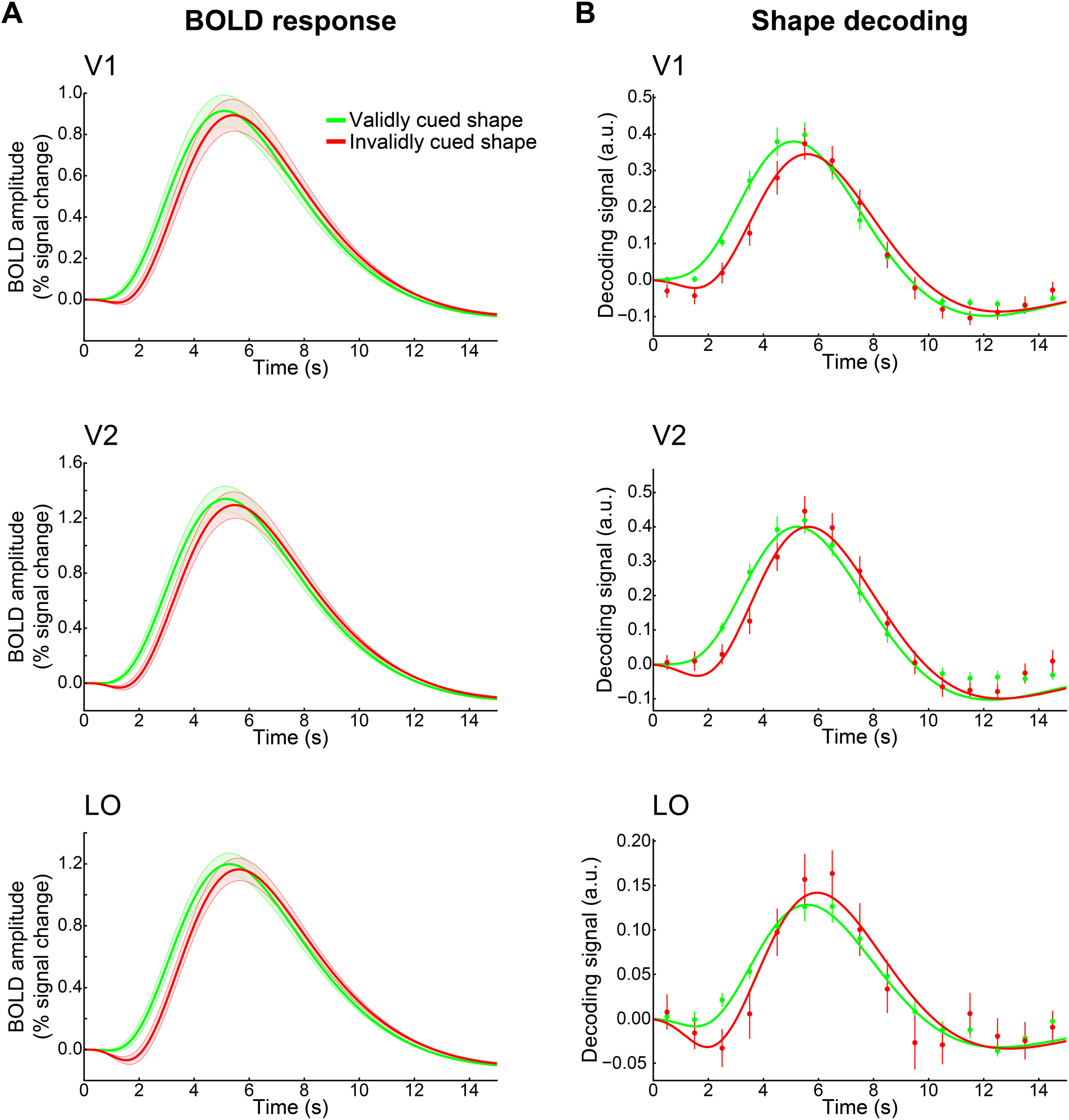
Time-resolved activity and decoding in visual cortex. (**A**) Timecourse of the mean BOLD response, separately for validly (green) and invalidly (red) predicted shapes, in visual cortex. These timecourses reflect the fit of the canonical HRF and its temporal derivative to the preprocessed fMRI data by condition. (**B**) Timecourse of the shape decoding signal, separately for validly (green) and invalidly (red) predicted shapes. Here, the canonical HRF and its derivative were not fit to the fMRI data directly, but rather to a continuous decoding signal obtained by reconstructing shape information for each timepoint from FIR parameter estimates. Shaded regions and error bars indicate SEM.

The delay for invalidly predicted shapes was also apparent in the temporal evolution of the reconstructed shape representations (Figure 4B). There was a reliable difference in the temporal derivative of the time-resolved decoding signal in V1 (*t*_23_ = 3.40, *p* = 0.0024) and V2 (*t*_23_ = 3.06, *p* = 0.0056), with a marginal effect in LO (*t*_23_ = 1.96, *p* = 0.062). In V1, the peak of the decoding signal was significantly lower for invalidly predicted shapes than for validly predicted shapes (*t*_23_ = 2.73, *p* = 0.012), while there was no such effect in V2 (*t*_23_ = 1.11, *p* = 0.27) or LO (*t*_23_ = 0.15, *p* = 0.87).

In sum, although there was a modest effect of prediction on the amplitude of the shape decoding signal in V1, the most striking effects of prediction in visual cortex were on the *latency* of the BOLD response and decoding signal.

### Hippocampal-cortical relationships

It is impossible with fMRI to establish that hippocampal prediction causes visual facilitation, but a precondition for such a mechanism is that these two measures should be related. Testing this relationship within participants was not possible in the current study because single-trial reconstruction and decoding of predictions was too noisy, especially in the hippocampus. We thus adopted an across-participant approach: We hypothesised that participants with greater decoding of the predicted shape in the hippocampus should have a greater latency shift in the decoding of invalidly vs. validly cued shapes in visual cortex. We found such a relationship between the hippocampus and LO (*r* = 0.42, *p* = 0.040; Figure 5), but not with V1 (*r* = −0.29, *p* = 0.17) or V2 (*r* = −0.05, *p* = 0.81).

**Figure 5.**
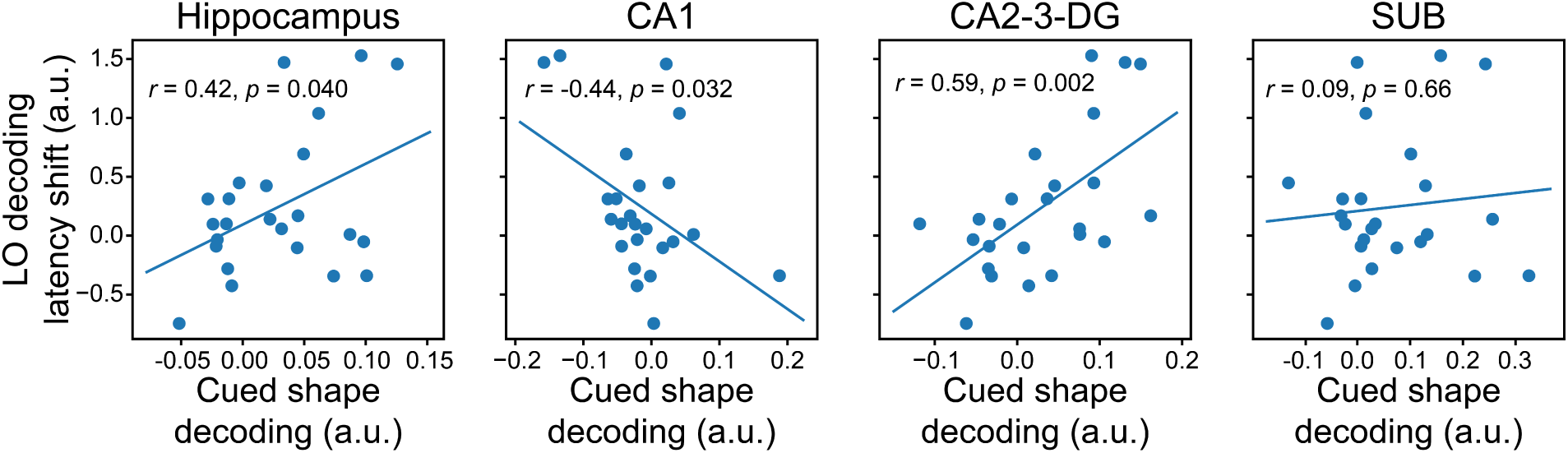
Hippocampal-cortical interactions. Correlation between the strength of predicted shape decoding in the hippocampus and the latency shift in the decoding signal caused by predictions in LO.

Strikingly, the hippocampal-LO relationship differed strongly across hippocampal subfields (Figure 5). In CA2-CA3-DG, as for the hippocampus as a whole, there was a reliable positive relationship (*r* = 0.59, *p* = 0.0022), with more hippocampal prediction associated with more LO facilitation. In CA1, however, the relationship was reliably *negative* (*r* = −0.44, *p* = 0.032), with more hippocampal prediction associated with *less* LO facilitation. There was no reliable relationship in subiculum (*r* = 0.09, *p* = 0.66). The surprising negative relationship between CA1 and LO also held for V1 (*r* = −0.55, *p* = 0.0047) and V2 (*r* = −0.57, *p* = 0.0036), whereas the positive relationship of CA2-CA3-DG was found only for LO (V1: *r* = 0.039, *p* = 0.86; V2: *r* = 0.17, *p* = 0.43).

In sum, although these findings do not resolve the causal direction of hippocampal-cortical interactions, they are consistent with the proposed mechanism of the hippocampus supplying predictions to visual cortex, compared to if we had found no such relationships.

### Caudate predictions

In addition to the hippocampus, we also examined the striatum, specifically the caudate and putamen, based on previous studies of prediction (Den Ouden et al., 2009; Turk-Browne et al., 2009), as well as the known involvement of the striatum in associative learning (Poldrack et al., 2001; Shohamy and Turk-Browne, 2013). We found that the caudate represented the predicted shape (*t*_23_ = 3.07, *p* = 0.0054) but not presented shape (*t*_23_ = 0.10, *p* = 0.92), as in the hippocampus. We could not reconstruct shape information from the putamen, for either the predicted (*t*_23_ = 1.58, *p* = 0.13) or presented (*t*_23_ = 0.27, *p* = 0.79) shapes. Unlike the hippocampus, LO facilitation did not correlate with prediction in the caudate (*r* = 0.23, *p* = 0.27) or putamen (*r* = 0.23, *p* = 0.27), nor were there correlations with V1 or V2 (*p*s > 0.05).

## Discussion

Predictive coding refers to a class of theories (Mumford, 1992; Rao and Ballard, 1999; Friston, 2005) about cortical processing that fits well with the strong influence of predictions on sensory processing reported here and elsewhere (Den Ouden et al., 2009; Alink et al., 2010; Meyer and Olson, 2011; Todorovic et al., 2011; Wacongne et al., 2011; Kok et al., 2012). Such models contain separate populations of neurons encoding predictions and prediction errors (Bell et al., 2016; Fiser et al., 2016; Kok, 2016; Kok et al., 2016a) and seek to explain mostly lower-level phenomena that can be resolved within local circuits of visual cortex, such as end-stopping and surround suppression (Rao and Ballard, 1999; Spratling, 2010). However, many predictive cues in our environment require cross-modal interactions and invoke complex expectations about objects. In these situations, the source of the prediction signals may lie outside of visual cortex, a possibility that has not received much attention in the predictive coding literature.

We hypothesised that such cross-modal predictions may be generated in the hippocampus. Specifically, upon presentation of a predictive cue, CA3 may retrieve the associated item through pattern completion of a learned temporal relation, and send this prediction to CA1, and from there back to sensory cortex, including through the subiculum (Lavenex and Amaral, 2000). Within CA1, memory-based predictions (originating from CA3) have been proposed to inhibit matching sensory signals (from EC), thereby signalling novelty, or prediction error (Lisman and Grace, 2005; Kumaran and Maguire, 2007; Chen et al., 2011; Duncan et al., 2012; Chen et al., 2015). Based on this model, one would expect CA3, but not CA1, to represent only the predicted item, and that is indeed what we observed in the current study. In addition, a representation of the predicted item was found in the subiculum, known to be a major relay between the hippocampus and sensory cortex, though admittedly not well understood and often excluded from hippocampal models (McClelland et al., 1995; Schapiro et al., 2017).

If the hippocampus is a source of sensory expectations, there should be a relationship between the strength of hippocampal predictions and effects of prediction in visual cortex. Although the current study did not allow us to study this relationship within participant (across trials) because of poor single-trial decoding, we did find such a relationship across participants. This also held for the CA2-CA3-DG subfield alone, in line with CA3’s proposed role in generating predictions via pattern completion. Conversely, in line with the proposed inhibitory role of predictions in CA1, prediction strength in this subfield was *negatively* correlated with the facilitative effects of prediction on visual cortex.

The model outlined above suggests a specific direction of neural signal flow during the generation of predictions, namely from CA3 through CA1 and the subiculum to cortex. However, due to the slow nature of the haemodynamic response and the lack of causal intervention, standard fMRI does not allow us to distinguish the direction of flow between regions. Future studies will be needed to directly address this important issue, including with intracranial recordings in neurological patients with both depth electrodes in the hippocampus and surface electrodes in sensory cortex. Additionally, signals from the hippocampus to cortex are known to arrive in the deep layers of EC, while signals from cortex to hippocampus flow through the superficial layers of EC (Lavenex and Amaral, 2000). Using high-field fMRI to study layer-specific prediction signals in EC could thus be used to help establish the direction of signal flow between hippocampus and cortex (Muckli et al., 2015; Kok et al., 2016a).

The current study suggests that the hippocampus is involved in signalling cross-modal predictions. However, there are several other mechanisms for prediction in the brain, including related to object recognition and semantic labels in medial prefrontal cortex (Bar et al., 2006) and to value and reinforcement learning in the ventral striatum (Den Ouden et al., 2012), as well as many other areas of polymodal association cortex that receive the required sensory inputs. What might distinguish the contribution of the hippocampus is the ability to quickly and flexibly learn new predictions, whereas these other systems learn more gradually after extensive experience and consolidation (McClelland et al., 1995; Schapiro et al., 2017). Regardless, further work will be needed to understand the relative contributions of each system and whether they have a cooperative or competitive relationship. For instance, it has long been unclear whether the ventral striatum contains stimulus-specific predictions. It has been proposed that the striatum serves as a gating mechanism, upregulating connectivity between top-down attention systems and sensory cortex when prediction errors occur (Zink et al., 2006; Den Ouden et al., 2010), rather than containing actual stimulus representations itself. However, our findings suggest that at least the caudate contains shape-specific representations of predicted stimuli and that they are encoded in similar activity patterns to the corresponding sensory stimuli (given generalisation of the model from localiser to prediction runs). An important avenue for future research would be to tease apart the roles of these two learning systems, the hippocampus and the striatum, in storing predictive associations, and their respective roles in sending feedback to sensory cortex (Poldrack et al., 2001; Shohamy and Turk-Browne, 2013).

The effects of the complex shape predictions on processing in the visual cortex, as reported here, differ strikingly from those reported previously for low-level feature predictions, using an otherwise similar paradigm (Kok et al., 2012). Whereas invalid grating orientation predictions in that study led to both an increased peak BOLD amplitude and a reduced orientation representation in V1 (Kok et al., 2012), the current study found that invalid shape predictions lead to *delayed* signals, both in terms of BOLD amplitude and shape representations. Although the cause of this difference is currently unclear, we offer a couple of potential explanations: First, predictions about low-level features and complex shapes may be encoded differently in visual cortex. Whereas a prediction about grating orientation could be encoded by simply increasing the gain of all neurons tuned for that orientation across the visual field (Kok et al., 2016b), complex shape predictions would require encoding different orientations and curvatures at specific retinotopic locations. This may be a particularly difficult challenge given that complex shapes are known to be encoded in a spatially invariant manner in higher-level visual cortex (DiCarlo et al., 2012). In line with this account, there is evidence that predictions about low-level features and complex natural images can have different effects on perception (Denison et al., 2011, 2016). Second, whereas invalid gratings in Kok et al. (2012) were maximally different from predicted gratings (i.e., orthogonal orientations), the difference between predicted and unpredicted shapes was more subtle. Such small violations may be less prone to strong prediction errors, but may rather lead to an *integration* of top-down predictions and bottom-up sensory signals (Kok et al., 2013). Clearly, future research is required to investigate these and other factors. A clear next step is to investigate how low-level feature predictions, particularly when involving cross-modal cues, engage the hippocampus.

How might the latency differences between visual cortex signals induced by validly and invalidly predicted shapes be explained, in terms of underlying neural signals? We are of course not suggesting that neuronal spiking in visual cortex is delayed half a second by invalid prediction cues. Rather, invalid predictions may lead to the suppression of visual cortex signals evoked by the first shape on a given trial, but less so for the second shape (which is no longer really unexpected once the first shape has been observed). If only the response to the first shape, but not the second shape, is suppressed, this could lead to a delayed peak activity once convolved with the BOLD response. This scenario seems particularly plausible for the reconstructed shape representations, since the early BOLD signals on invalid trials would presumably contain a mixture of the predicted and presented shapes, which might to some extent cancel each other out in the eyes of the decoder.

Previous studies have found that predictive cues can lead to the cortical reinstatement of expected stimuli (Kok et al., 2014; Hindy et al., 2016), in anticipation of the actual sensory inputs (Kok et al., 2017). Such cortical reinstatement has also been shown for other cognitive processes as well, such as visual short term memory (Harrison and Tong, 2009; Bosch et al., 2014; Gordon et al., 2014), mental imagery (Stokes et al., 2009a; Albers et al., 2013), and preparatory attention (Stokes et al., 2009b; Peelen and Kastner, 2011; Myers et al., 2015). One intriguing possibility is that these different cognitive processes are subserved by the same neural mechanism (Pearson and Westbrook, 2015). Specifically, is cortical reinstatement in working memory, imagery, and attention mediated by the hippocampus, as it seems to be in associative memory (Bosch et al., 2014; Gordon et al., 2014) and the cross-modal predictions studied here? Additionally, regardless of the source, do these different processes affect visual cortex the same way, or do different processes modulate different layers of visual cortex, in support of different computational goals (Friston, 2005; Muckli et al., 2015; Kok et al., 2016a)?

In conclusion, here we find that patterns of neural activity in the hippocampus reflect stimulus-specific predictions, as signalled by cross-modal cues. These findings are consistent with a role for the hippocampus in generating memory-based predictions that influence sensory systems.

## Materials and Methods

### Participants

Twenty-five healthy individuals participated in the experiment. All participants were right-handed, MR-compatible, and had normal or corrected-to-normal vision. Participants provided informed consent to a protocol approved by the Princeton University Institutional Review Board and were compensated ($20 per hour). One participant was excluded from analysis because they moved their head between runs such that large parts of the occipital lobe were no longer inside the field of view. The final sample consisted of 24 participants (15 female, age 23 ± 3, mean ± SD).

### Stimuli

Visual stimuli were generated using MATLAB (Mathworks, Natick, MA, USA) and the Psychophysics Toolbox (Brainard, 1997). In the MR scanner, the stimuli were displayed on a rear-projection screen using a projector (1024 × 768 resolution, 60 Hz refresh rate) against a uniform grey background. Participants viewed the visual display through a mirror that was mounted on the head coil. The visual stimuli consisted of complex shapes defined radial frequency components (RFCs) (Zahn and Roskies, 1972; Op de Beeck et al., 2001; Drucker and Aguirre, 2009) (Figure 1A). The contours of the stimuli were defined by seven RFCs, based on a subset of the stimuli used in Op de Beeck et al. (2001) (see their Figure 1a). A one-dimensional shape space was created by varying the amplitude of three out of the seven RFCs. Specifically, the amplitudes of the 1.11 Hz, 1.54 Hz and 4.94 Hz components increased together, ranging from 0 to 36 (first two components), and from 15.58 to 33.58 (third component). Note that we chose to vary three RFCs simultaneously, rather than one, to increase the perceptual (and neural) discriminability of the shapes.

In order to map out this shape space perceptually, we generated 13 shapes that spanned the continuum, with the amplitudes of the three modulated RFC increasing with equal steps from the minimum to the maximum of the ranges defined above. Six participants categorised these shapes as one of the two extremes of the continuum (each shape presented 24 times). Psychometric curves were fit to these data and we determined the points along the continuum (in terms of the amplitudes of the three modulated RFCs) that were judged as 10%, 50% and 90% likely to be the extreme shape with maximal values. The five shapes we used in the fMRI experiment consisted of these three experimentally determined points in the space continuum, as well as the two extremes (Figure 1A). The participants who took part in the fMRI experiment were exposed to the same perceptual categorisation experiment after the fMRI session ended, and we determined that for each participant the chance of classifying a shape as the maximal extreme increased monotonically as a function of the amplitude of the three RFCs. During the fMRI experiment, the shapes were presented centred on fixation (colour: black, size: 4.5°). Additionally, a fourth RFC (the 3.18 Hz component) was used to create slightly warped versions of the five shapes, to enable the same/different shape discrimination cover task (see below).

Auditory cues consisted of three pure tones (440, 554 and 659 Hz, 80 ms per tone, 5 ms intervals), presented in either ascending or descending pitch.

### Experimental design

Each trial of the main experiment started with the presentation of a fixation bullseye (diameter: 0.7°). During the prediction runs, an auditory cue (ascending or descending tones, 250 ms) was presented 100 ms after onset of the trial (Figure 1A). After a 500 ms delay two consecutive shape stimuli were presented for 250 ms each, separated by a 500 ms blank screen (Figure 1A). The auditory cue (ascending vs. descending tones) predicted whether the first shape on that trial would be shape 2 or shape 4 (out of five shapes; Figure 1B). The cue was valid on 75% of trials, while in the other 25% of trials the unpredicted shape would be presented. For instance, an ascending auditory cue might be followed by shape 2 on 75% of trials, and by shape 4 on the remaining 25% of trials. The contingencies between the auditory cues and grating orientations were flipped halfway through the experiment, and the order of mappings was counterbalanced across participants.

On each trial, the second shape was either identical to the first, or was slightly warped. This warp was achieved by modulating the amplitude of the 3.18 Hz RFC component defining the shape. This modulation could be either positive or negative (counterbalanced over conditions) and participants’ task was to indicate whether the two shapes on a given trial were the same or different, using an MR-compatible button box. After the response interval ended (750 ms after disappearance of the second shape), the fixation bullseye was replaced by a single dot, signalling the end of the trial while still requiring participants to fixate. This task was designed to avoid a direct relationship between the perceptual prediction and the task response. Furthermore, by modulating one of the RFCs that was not used to define our one-dimensional shape space, we ensured that the shape change on which the task was performed was orthogonal to the changes that defined the shape space, and thus orthogonal to the prediction cues. The size of the modulation was determined by an adaptive staircasing procedure (Watson and Pelli, 1983), updated after each trial, in order to make the task challenging (~75% correct). Separate staircases were run for trials containing valid and invalid cues, as well as for the localiser runs, to equate task difficulty between conditions. All participants completed two runs of this task (128 trials per run).

In two additional runs, which were interleaved with the runs just described (in ABBA fashion, order counterbalanced over participants), the 25% invalid trials did not involve presentation of the unpredicted shape, but rather no shape stimuli were presented at all. These omission trials were included in an attempt to decode expected but omitted shapes from the BOLD response. However, no such effects were found. We are conducting additional studies to better understand the conditions under which omission trials do (e.g., Kok et al., 2014; Hindy et al., 2016) and do not (the current study) reveal expectations, and thus the data from the omission trials will not be considered further in this manuscript.

Finally, participants completed two localiser runs (120 trials per run), in which no auditory cues were presented. As in the prediction runs, the start of each trial was signalled by the onset of the fixation bullseye, and the SOA between this onset and the presentation of the first shape was 850 ms (Figure 1C). On any given trial, one of the five shapes would be presented, with equal (20%) likelihood. As in the prediction runs, the first shape was followed by a second one that was either identical or slightly warped, and participants’ task was to report same or different. These runs were designed to be as similar as possible to the prediction runs, save the absence of the auditory cues and the equal rates of presentation of all five shapes. The two localiser runs flanked the runs containing the auditory cues, constituting the first run and sixth (last) run of the experiment.

The staircases were kept running throughout the experiment. They were initialised at a value determined during a practice session 1-3 days before the fMRI experiment (no auditory cues, 120 trials). After the initial practice run, the meaning of the auditory cues was explained, and participants practiced briefly with both cue-shape contingencies (valid trials only; 16 trials per contingency). Additionally, before the first prediction run of the fMRI experiment (run 2), as well as in between runs 3 and 4, when the contingencies between cue and stimuli were flipped, participants performed a practice block (112 trials, valid cues only), to acquaint them with the upcoming cue-shape contingency.

### MRI acquisition

Structural and functional MRI data were collected on a 3T Siemens Prisma scanner with a 64-channel head coil. Functional images were acquired using a multiband echo-planar imaging sequence (TR = 1000 ms, TE= 32.6 ms, 60 transversal slices, voxel size = 1.5 × 1.5 × 1.5 mm, 55° flip angle, multiband factor 6). This sequence produced a partial volume for each participant, parallel to the hippocampus and covering the majority of the temporal and occipital lobes. Anatomical images were acquired using a T1-weighted MPRAGE sequence, using a GRAPPA acceleration factor of 3 (TR = 2300 ms, TE = 2.27 ms, voxel size 1 × 1 × 1 mm, 192 transversal slices, 8° flip angle). Additionally, to enable hippocampal segmentation, two T2-weighted turbo spin-echo (TSE) images (TR = 11390 ms, TE = 90 ms, voxel size = 0.44 × 0.44 × 1.5 mm, 54 coronal slices, perpendicular to the long axis of the hippocampus, distance factor 20%, 150° flip angle) were acquired. To correct for susceptibility-induced distortions in the EPI images, a pair of spin echo volumes was acquired in opposing phase encode directions (anterior/posterior and posterior/anterior) with matching slice prescription, voxel size, field of view, bandwidth, and echo spacing (TR = 8000 ms, TE = 66 ms).

### fMRI preprocessing

The images were preprocessed using FEAT 6 (FMRI Expert Analysis Tool), part of FSL 5 (http://fsl.fmrib.ox.ac.uk/fsl, Oxford Centre for Functional MRI of the Brain) (Jenkinson et al., 2012). Susceptibility-induced distortions were determined on the basis of the opposing spin echo volume pairs using the FSL topup tool (Andersson et al., 2003). The resulting off-resonance field output was converted from Hz to rad/s, and supplied to FEAT for B0 unwarping (see below). The first six volumes of each run were discarded to allow T1 equilibration. For each run, the remaining functional images were spatially realigned to correct for head motion, and simultaneously supplied to B0 unwarping and registered to the participants’ structural T1 image, using boundary-based registration. The functional data were temporally high-pass filtered with a 128 s period cutoff, no spatial smoothing was applied. Finally, the two TSE images were averaged and the resulting image was registered to the T1 image through FMRIB’s Linear Image Registration Tool (FLIRT).

All analyses were performed in participants’ native space. For the searchlight analyses (see below), each participant’s output volumes were registered to the MNI (Montreal Neurological Institute) template to allow group-level statistics. This was achieved by applying the non-linear registration parameters obtained from registering each participant’s T1 image to the MNI template using AFNI’s 3dQwarp (https://afni.nimh.nih.gov/pub/dist/doc/program_help/3dQwarp.html).

### Regions of interest

Hippocampal subfields CA2–CA3–DG, CA1, and the subiculum, were defined on the basis of the TSE and T1 images using the automatic segmentation of hippocampal subfields (ASHS) machine learning toolbox (Yushkevich et al., 2015) and a database of manual medial temporal lobe (MTL) segmentations from a separate set of 51 participants (Aly and Turk-Browne, 2016a, 2016b). Manual segmentations were based on anatomical landmarks used in prior studies (Duvernoy, 2005; Carr et al., 2010; Schapiro et al., 2012). Consistent with these studies, CA2, CA3 and DG were combined into a single ROI because these subfields are difficult to distinguish at our functional resolution (1.5 mm isotropic). TSE acquisition failed for one participant, and so their hippocampal ROIs were based on the T1 image alone. Results of the automated segmentation were inspected visually for each participant. The hippocampus ROI consisted of the union of the CA2–CA3–DG, CA1, and subiculum subfields.

In visual cortex, V1, V2 and lateral occipital (LO) cortex automatically defined in each participant’s T1-weighted anatomical scan with FreeSurfer (http://surfer.nmr.mgh.harvard.edu/). Finally, putamen and caudate ROIs were obtained from FreeSurfer’s subcortical segmentation, since these regions have been implicated in associative learning and prediction (Poldrack et al., 2001; Den Ouden et al., 2009; Turk-Browne et al., 2009; Shohamy and Turk-Browne, 2013).

The visual cortex ROIs were restricted to the 500 most active voxels during the localiser runs in each ROI, to ensure that we were measuring responses in the retinotopic locations corresponding to our visual stimuli. Since no clear retinotopic organization is present in the other ROIs, cross-validated feature selection was used instead (see below).

All ROIs were collapsed over the left and right hemispheres, since we had no hypotheses regarding hemispheric differences.

### fMRI data modelling

The functional data of each participant were modelled with general linear model (GLM), using FMRIB’s Improved Linear Model (FILM), which included temporal autocorrelation correction and extended motion parameters (6 standard parameters, plus their derivatives and their squares) as nuisance covariates. We specified regressors for the conditions of interest (localiser runs: five shapes; prediction runs: two shapes x two prediction conditions [valid vs. invalid]), by convolving a delta function at the onset of the first shape on each trial with a double-gamma HRF. Additionally, we included the temporal derivative of each regressor in order to accommodate variability in the onset of the response (Friston et al., 1998).

To investigate the temporal evolution of shape representations in visual cortex, a finite impulse response (FIR) approach was used to estimate the BOLD signal evoked by each condition of interest in 20 1s intervals. This allowed us to estimate the shape decoding signal in a time-resolved manner, by training the decoder on the FIR parameter estimates from the 4-7s time bins in the localiser runs (corresponding to the peak hemodynamic signal) and applying it to all time bins for the prediction runs. The amplitude and latency of this time-resolved decoding signal was quantified by fitting a double-gamma function and its temporal derivative.

### Shape decoding

In order to probe neural shape representations, we used a forward modelling approach to reconstruct the shape from the pattern of BOLD activity in a given brain region (Brouwer and Heeger, 2009). This approach has proven successful in reconstructing continuous stimulus features, such as hue (Brouwer and Heeger, 2009), orientation (Brouwer and Heeger, 2011), and motion direction (Kok et al., 2013). In the current study, shape contour was constructed along a continuous dimension (see above), allowing the application of a forward model.

We characterised the shape selectivity of each voxel as a weighted sum of five hypothetical channels, each with an idealised shape tuning curve (or basis function). As in previous forward model implementations (Brouwer and Heeger, 2009, 2011; Kok et al., 2013), each basis function consisted of a halfwave-rectified sinusoid raised to the fifth power, and the five basis functions were spaced evenly, such that they were centred on the five points in shape space that constituted the five shapes presented in the experiment (Figure 2A). As a result of this, a tuning curve with any possible shape preference (within the space defined here) could be expressed as a weighted sum of the five basis functions. Note that, unlike other stimulus features previously reconstructed using forward models, the shape space used here was not circular, and therefore the channels did not wrap around.

In the first stage of the analysis, we used parameter estimates obtained from the two localiser runs to estimate the weights on the five hypothetical channels separately for each voxel, using linear regression. Specifically, let *k* be the number of channels, *m* the number of voxels, and *n* the number of measurements (i.e., the five shapes). The matrix of estimated response amplitudes for the different shapes during the localiser runs (**B**_**loc**_, *m × n*) was related to the matrix of hypothetical channel outputs (**C**_**loc**_, *k × n*) by a weight matrix (**W***, m × k*):

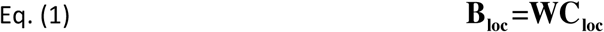

The least-squares estimate of this weight matrix **W** was estimated using linear regression:

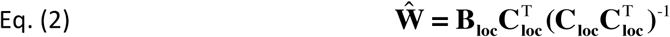

These weights reflected the relative contribution of the five hypothetical channels in the forward model (each with their own shape selectivity) to the observed response amplitude of each voxel. Using these weights, the second stage of analysis reconstructed the channel outputs associated with the pattern of activity across voxels evoked by the stimuli in the main experiment (**B**_**exp**_), again using linear regression. This step transformed each vector of *n* voxel responses (parameter estimates per condition) into a vector of 5 (number of basis functions) channel responses. More specifically, the channel responses (**C**_**exp**_) associated with the responses in the main experiment (**B**_**exp**_) were estimated using the learned weights (**W**):

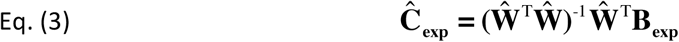

These channel outputs were used to compute a weighted average of the five basis functions, reflecting a neural shape tuning curve (Figure 2B). Note that, during the main experiment (i.e., the prediction runs), only shapes 2 and 4 were presented. Thus, four shape tuning curves were obtained for the prediction runs: two shapes by two prediction conditions (valid vs. invalid). We collapsed across the presented shape by subtracting the shape tuning curve for shape 4 from that for shape 2, thereby subtracting out any non-shape-specific BOLD signals (Figure 2C).

Decoding performance was quantified by subtracting the amplitude of the shape tuning curve at the presented shape (e.g., shape 2) from the amplitude at the non-presented shape (shape 4). Collapsing across conditions led to two measures of decoder evidence per participant: one for validly predicted shapes, and one for invalidly predicted shapes. This allowed us to quantify evidence for the shape as presented on the screen (by averaging evidence for validly and invalidly predicted shapes) and evidence for the cued shape (by averaging (1 - evidence) for the invalidly predicted shapes with evidence for the validly predicted shapes). These measures were statistically tested at the group level using simple t-tests.

For the visual cortex ROIs, voxels were selected based on the strength of the evoked response to the shapes during the localiser runs. Other brain regions, such as the hippocampus, do not show a clear evoked response to visual stimuli. Therefore, we followed a different voxel selection procedure for the other ROIs. First, voxels were sorted by their informativeness, that is, how different the weights for the five channels were from each other (quantified by the standard deviation of the five weights). Second, the number of voxels to include was determined by selecting between 10% and 100% of all voxels (in steps of 10%), and training and testing the model on these voxels, within the localiser runs (i.e., training on one run and testing on the other run). For each iteration, decoding performance on shapes 2 and 4 was quantified as described above, and the number of voxels that yielded the highest decoding performance was selected (group average: hippocampus, 1536 of 3383 voxels; caudate, 590 of 2240 voxels; putamen, 1498 of 3582 voxels).

We also labelled the selected hippocampus voxels based on their subfield from the hippocampal segmentation (group average: CA1, 436 voxels; CA2-CA3-DG, 572 voxels; subiculum, 425 voxels). Differential contributions of the subfields were statistically tested by performing a one-way repeated measures ANOVA on the measure of interest (e.g., decoding of the cued shape; Figure 3D).

For the main ROI and searchlight analyses, the input to the forward model consisted of voxelwise double-gamma parameter estimates, reflecting the amplitude of the BOLD response. Additionally, decoding was also applied to the FIR model parameter estimates in visual cortex.

### Searchlight analysis

In order to explore the specificity of presented and predicted shape representations, a multivariate searchlight approach was used to test these effects within the field of view of our functional scans (most of occipital and temporal, and part of parietal and frontal cortex). A spherical searchlight with a radius of 5 voxels (7.5 mm) was passed over all functional voxels. In each searchlight, we performed shape decoding in the same manner as in the ROIs, yielding maps of decoder evidence for the presented and predicted shapes, respectively, for each participant. Group-level non-parametric permutation tests were applied to these searchlight maps using FSL Randomise (Winkler et al., 2014), correcting for multiple comparisons at *p* < 0.05 using threshold-free cluster enhancement (Smith and Nichols, 2009).

## Acknowledgements

This work was supported by an NWO Rubicon grant 446-15-004 to P.K. and NIH R01 EY021755 and NIH R01 MH069456 to N.B.T.-B. The authors would like to thank Nicholas C Hindy and Mariam Aly for help with hippocampal segmentations.

## Competing interests

N.B.T.-B.: Reviewing editor, eLife. P.K. declares no competing interests.

